# Multi-regional Neurodegeneration in Alzheimer’s disease: Meta-analysis and data integration of transcriptomics data

**DOI:** 10.1101/245571

**Authors:** Karbalaei Reza, Rezaei-Tavirani Mostafa, Torkzaban Bahareh, Azimzadeh Sadegh

## Abstract

Alzheimer’s disease (AD) is a complex neurodegenerative disease with various deleterious perturbations in regulatory pathways of various brain regions. Thus, it would be critical to understanding the role of different regions of the brain in initiation and progression of AD, However, owing to complex and multifactorial nature of this disease, the molecular mechanism of AD has yet to be fully elucidated. To confront with this challenge, we launched a meta-analytical study of current transcriptomics data in four different regions of the brain in AD (Entorhinal, Hippocampus, Temporal and Frontal) with systems analysis of identifying involved signaling and metabolic pathways. We found different regulatory patterns in Entorhinal and Hippocampus regions to be associated with progression of AD. We also identified shared versus unique biological pathways and critical proteins among different brain regions. ACACB, GAPDH, ACLY, and EGFR were the most important proteins in Entorhinal, Frontal, Hippocampus and Temporal regions, respectively. Moreover, eight proteins including CDK5, ATP5G1, DNM1, GNG3, AP2M1, ALDOA, GPI, and TPI1 were differentially expressed in all four brain regions, among which, CDK5 and ATP5G1 were enriched in KEGG Alzheimer’s disease pathway as well.

## 1- Introduction

As a chronic neurodegenerative disorder, Alzheimer’s disease (AD) is exhibiting deleterious effects on patients, caregivers, and families [1]. Based on Alzheimer’s Disease International Federation (ADI) report at the current rate, 46.8 million people living with dementia worldwide, with numbers projected to nearly double every 20 years, increasing to 74.7 million by 2030 and 131.5 million by 2050 [2]. Also, they estimated that the Medicare/Medicaid spending in 2015 totaled US$818 billion (about four times more than what was predicted in 2005 namely US $216 billion) and estimates that the global cost of dementia will have reached US$1 trillion in 2018 [2].

Early AD diagnosis in combination with new classes of neuroprotective or disease-modifying drug treatments may delay or prevent the neurodegenerative effects of AD [3]. Attempts at symptomatic relief are only modestly effective, and still, there isn’t a proper report on some curative treatment [4], although many researchers focused on different aspects of this disease and tried to shed light on its diagnosis [5]. However, due to long preclinical and prodromal phases and the symptom-free episodes, the initiating factors of AD are still unclear, and it seems likely that a sort of brain damage starts a decade or more before problems become evident [5]. So, AD continuously is a major target of both clinical and basic types of research.

It is assumed that many factors and their interactions contribute to the pathogenesis of AD. Thus, a holistic research regarding the mechanism of AD is of great importance [6, 7]. AD is a multi-factor disease that has two forms, early onset familial Alzheimer disease (EFAD) that inherited in an autosomal dominant manner [8] and Late-onset Alzheimer’s disease (LOAD) or non-familial [9] that appears as a much more complex or multifactorial disease [10]. The point is that regardless the Alzheimer’s form there isn’t any significant diagnostic report before the appearance of Alzheimer’s hallmarks.

The AD’s hallmarks associated with microscopic features including appearance of Neurofibrillary Tangles [11], Amyloid Plaques [12], Cerebral Amyloid Angiopathy [13], Granulovacuolar Degeneration and Hirano Bodies [14], Glial Responses [15], Neuronal Loss [16] and Synapse Loss [17] which lead to appearance of the macroscopic features described by Braak in 1991 [18]. Braak stages of I and II represent the situation that neurofibrillary tangles of the transentorhinal region of the brain are involved. At the stages of III and IV the limbic region (such as the Hippocampus) is also involved and finally, at the stages of V and VI, an extensive neocortical involvement is detectable [18]. Altogether, abnormal deposits of proteins form amyloid plaques and tau tangles throughout the brain, leading to deficits in neurons plasticity and apoptosis and the resultant brain shrinkage over the time. With damage reaching the Hippocampus, (stage III and IV), substantial memory loss would be expected has been shrunken significantly [9].

Classical reductionist research methods are incapable of coping the complex nature of multi-factorial diseases such as AD in which, both individual’s genetic background and environmental conditions are involved. Thus a systems level methodology with an integrative and holistic approach would be highly demanding in this area to identify critical interactions between hereditary and environmental factors. Also, models were designed by these approaches to understanding initiation. To reach this goal, A large number of microarray datasets belonging to different brain regions of AD patients are available [19–21] in public databases. Herein, we used these data to shed light on the role of important pathways in brain regions associated with AD. In this regard we meta-analyzed qualified microarray data belonging to four brain regions of patients with AD including Entorhinal, Frontal, Hippocampus, and Temporal) and followed a network-based approach to identify brain-region-specific pathways/genes. Furthermore, we supplemented our results with current knowledge-and data-driven networks to decipher possible crosstalk between Hippocampus-Frontal in the brain regions of Alzheimer’s patients.

## 2- Materials and methods

### 2-1 Data collection

In general, 110 data-series belonging to 8 regions of brain deposited in GEO and Array Express databases based on “Alzheimer” and “Homo sapiens” keywords. To decrease heterogeneity as well as increasing the consistency between expressing data-series, only Human Genome U133 Plus 2.0 Array (HG-U133_Plus_2) or Human Genome U133A ver2.0 (U133A) platforms from Affymetrix Company were selected. Finally, the data-series of four regions including Entorhinal, Frontal, Hippocampus and Temporal out of eight brain regions were selected after quality assessment. A summary of the above procedures and the applied methods is presented in figure 1.

**Figure 1-.**
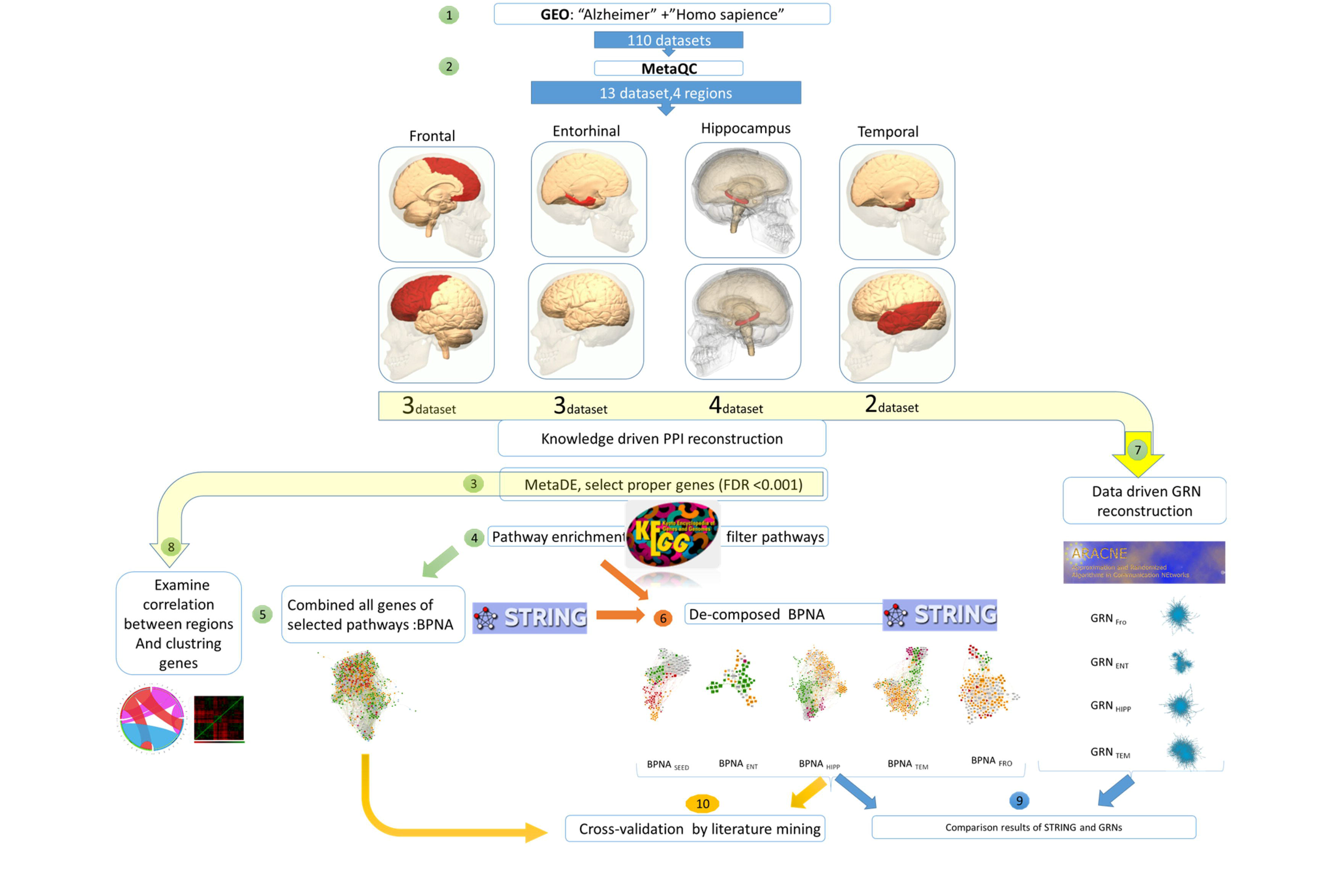
The Summary of the overall procedure and applied methods in this article. After selection and qualification proper Alzheimer datasets (step 1 and 2), MetaDE was performed on selected datasets (step 3) and DEGs were used for knowledge-driven analysis. First, pathways enrichment using KEGG was accomplished (step 4), then by combining all proteins of selected pathways (P-value < 0.05), BPNA network was constructed (step 5). By decomposing BPNA, five networks were reconstructed (step6). On the other side, on the best dataset in each region, GRN construction (ARACNe analysis) was performed (step 7), and pairwise correlation test was performed on all selected proteins from step 3 between regions (step 8). Results from step 6 compared with step 7 (step 9). Also, expression Condition of all proteins of BPNA (Up-regulated, Down-regulated or others) were validated in literature mining and compare with results of the meta-analysis were compared (step10). For more details, see materials and methods.

### 2-2 Gene expression data analysis and meta-analysis

To compile expression data for meta-analysis, CEL raw files were reprocessed in the following steps. First, quality of each microarray sample was assessed using the R package “array Quality Metrics” [22]. Second, normalization of raw data series was carried out using Just-RMA algorithm [23]. Third, expression data of each region of the brain was qualified for meta-analysis using MetaOmics package in R [24]. MetaOmics” software is a unified R package including several tools such as MetaQC and MetaDE. The MetaQC package performs principal component analysis (PCA) and standardized mean rank summary (SMR) score based on calculated six quality control (QC) measures to identify problematic studies and exclusion of heterogeneous samples [25] (Figure 2). Consequently, datasets for each brain region (Entorhinal, Frontal, Hippocampus, and Temporal) were qualified (Table 1). Afterward, a meta-analysis was performed using Fisher’s combined probability test in MetaDE package for each brain region [26] (Table S1). Benjamini-Hochberg Adjustment used for adjustment of p-values [27, 28]. Also, fold change of ≥I2I and false discovery rate (“FDR) ≤10−3” was used for identification of differential gene expression. In order to compare four brain regions, unique DEGs of each region (DEGs that only express in one region) was selected, sorted based on their fold change, and top 10% of DEGs were enriched for GO Biological Process (Table S2).

**Figure 2-.**
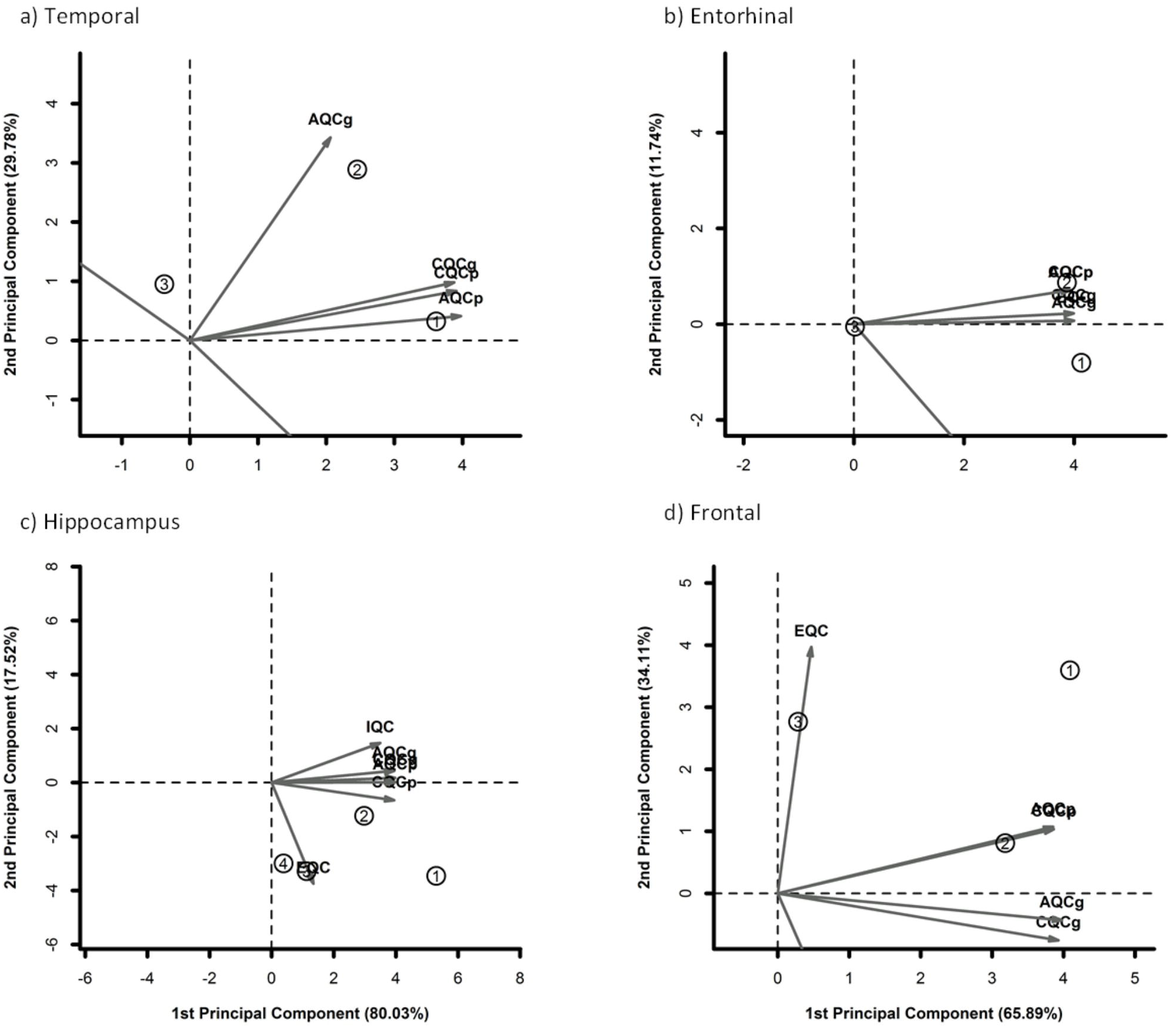
PCA biplots of meta-analysis. Studies placed opposite to quality axes are excluded. In the Temporal region, study GSE29652 was identified as an outlier and removed from the list (a)Temporal, b) Entorhinal, c) Hippocampus, and d)Frontal.

**Table 1-.**
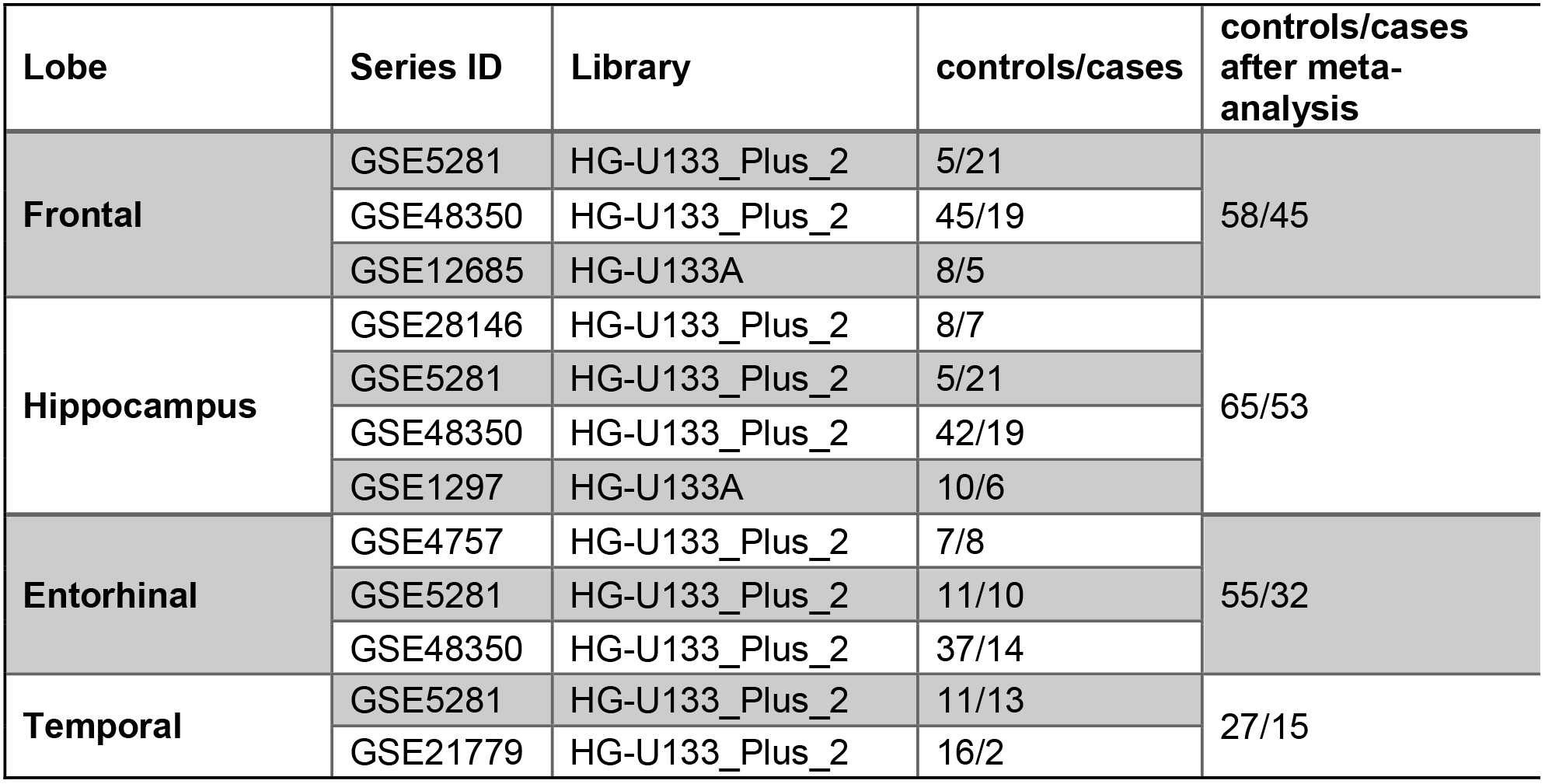
Selected datasets for Entorhinal, Frontal, Hippocampus and Temporal regions.

| Lobe | Series ID | Library | controls/cases | controls/cases after meta-analysis |
| --- | --- | --- | --- | --- |
| Frontal | GSE5281 | HG-U133_Plus_2 | 5/21 | 58/45 |
|  | GSE48350 | HG-U133_Plus_2 | 45/19 |  |
|  | GSE12685 | HG-U133A | 8/5 |  |
| Hippocampus | GSE28146 | HG-U133_Plus_2 | 8/7 | 65/53 |
|  | GSE5281 | HG-U133_Plus_2 | 5/21 |  |
|  | GSE48350 | HG-U133_Plus_2 | 42/19 |  |
|  | GSE1297 | HG-U133A | 10/6 |  |
| Entorhinal | GSE4757 | HG-U133_Plus_2 | 7/8 | 55/32 |
|  | GSE5281 | HG-U133_Plus_2 | 11/10 |  |
|  | GSE48350 | HG-U133_Plus_2 | 37/14 |  |
| Temporal | GSE5281 | HG-U133_Plus_2 | 11/13 | 27/15 |
|  | GSE21779 | HG-U133_Plus_2 | 16/2 |  |

### 2-3 Pathway enrichment and integration

The contribution of DEGs in different pathways was investigated using enrichment of DEGs in Enrichr web server [29, 30]. Enrichr performs enrichment analysis for the input genes against several data-libraries including complete data of Kyoto Encyclopedia of Genes and Genomes (KEGG) database (Release 78.0, April 1, 2016) [14, 31]. All significant pathways (p-value < 0.05) were identified for each meta-analyzed region of the brain involved in AD (Table 2). The enriched pathways were classified as “identical” assuming shared pathways among all four studied brain regions and “non-Identical” pathways, to address to other pathways which may be identical between two or three of them. Also, enriched pathways were categorized in three classes including signaling, metabolic, and disease (Table 2).

**Table 2-.**
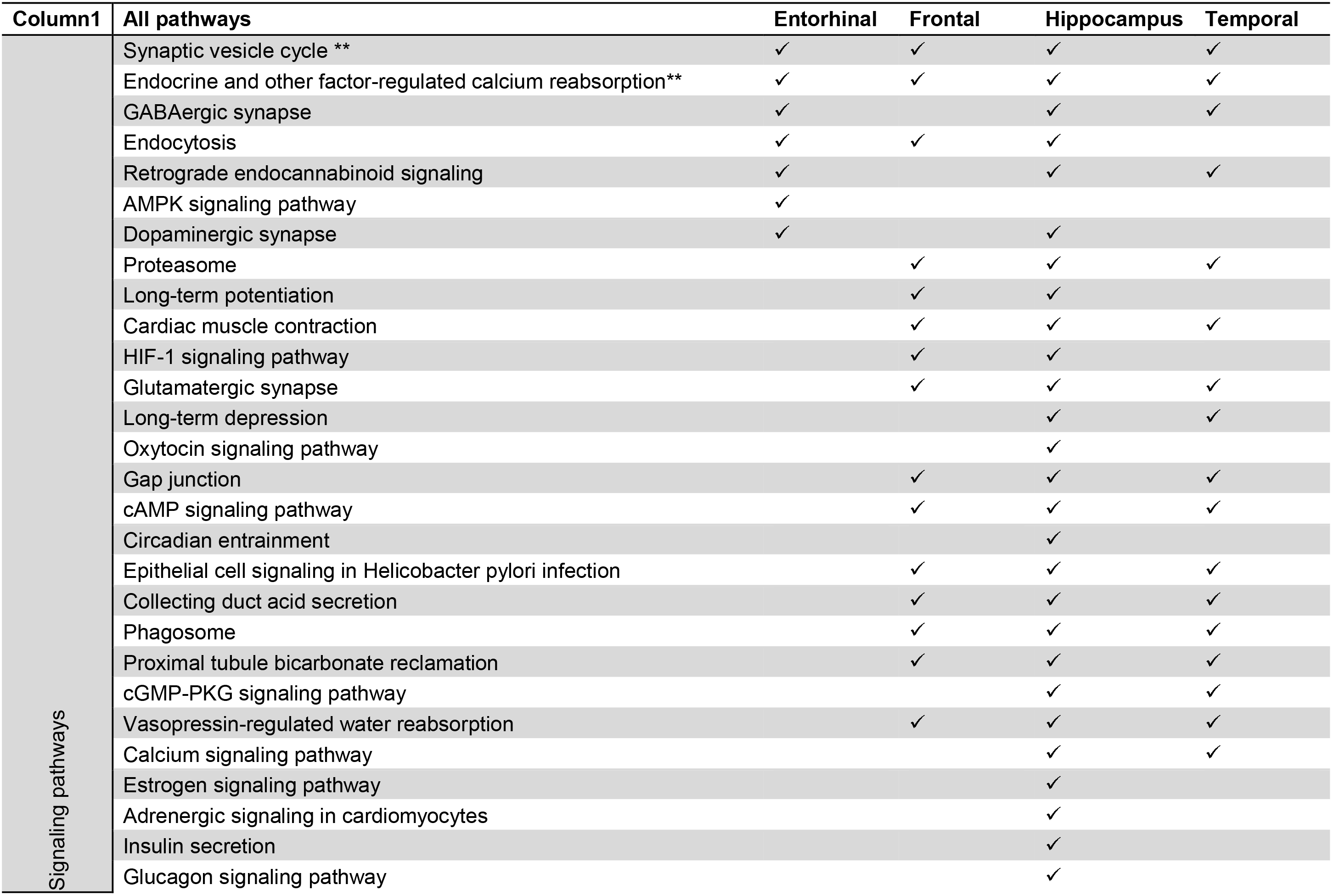
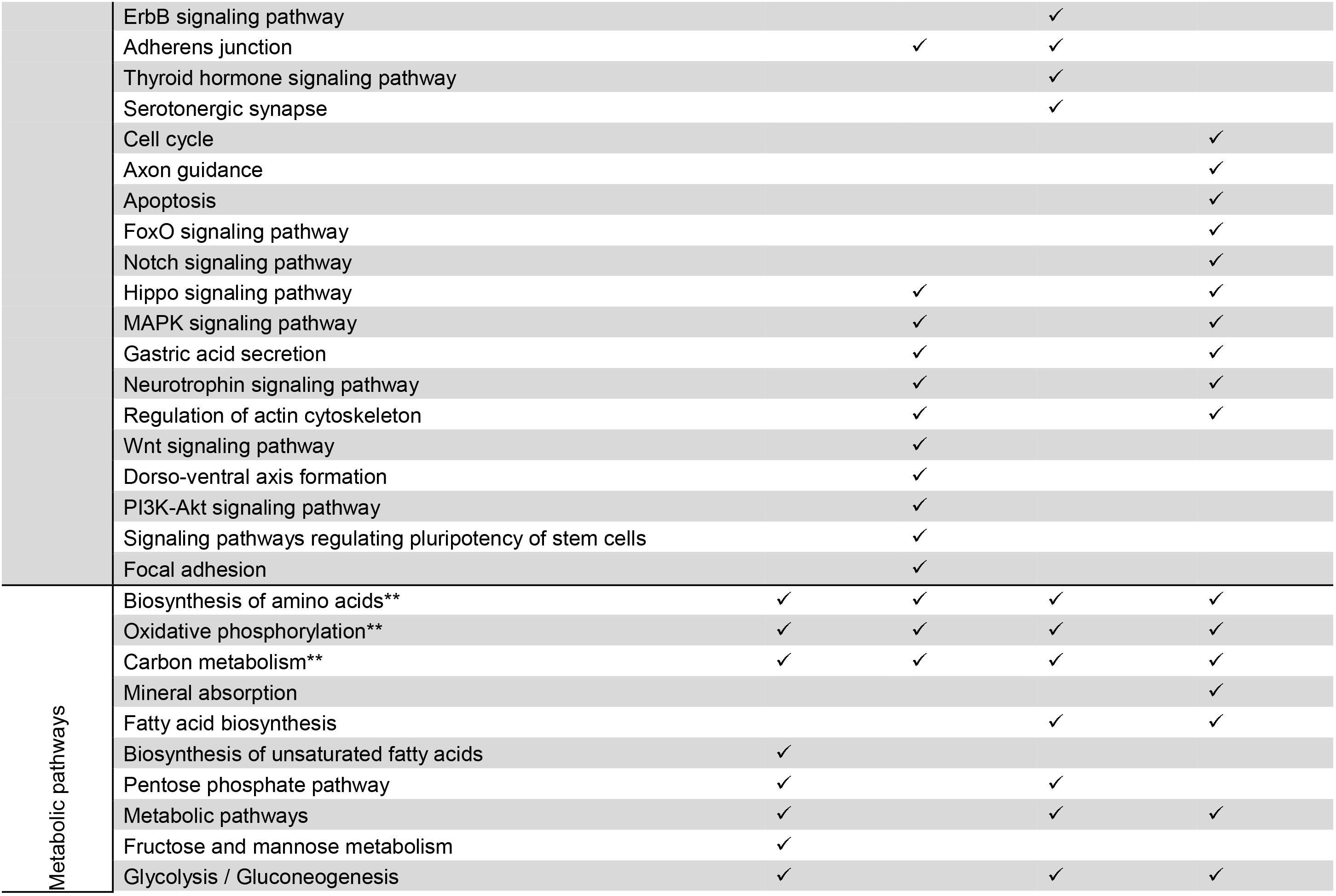
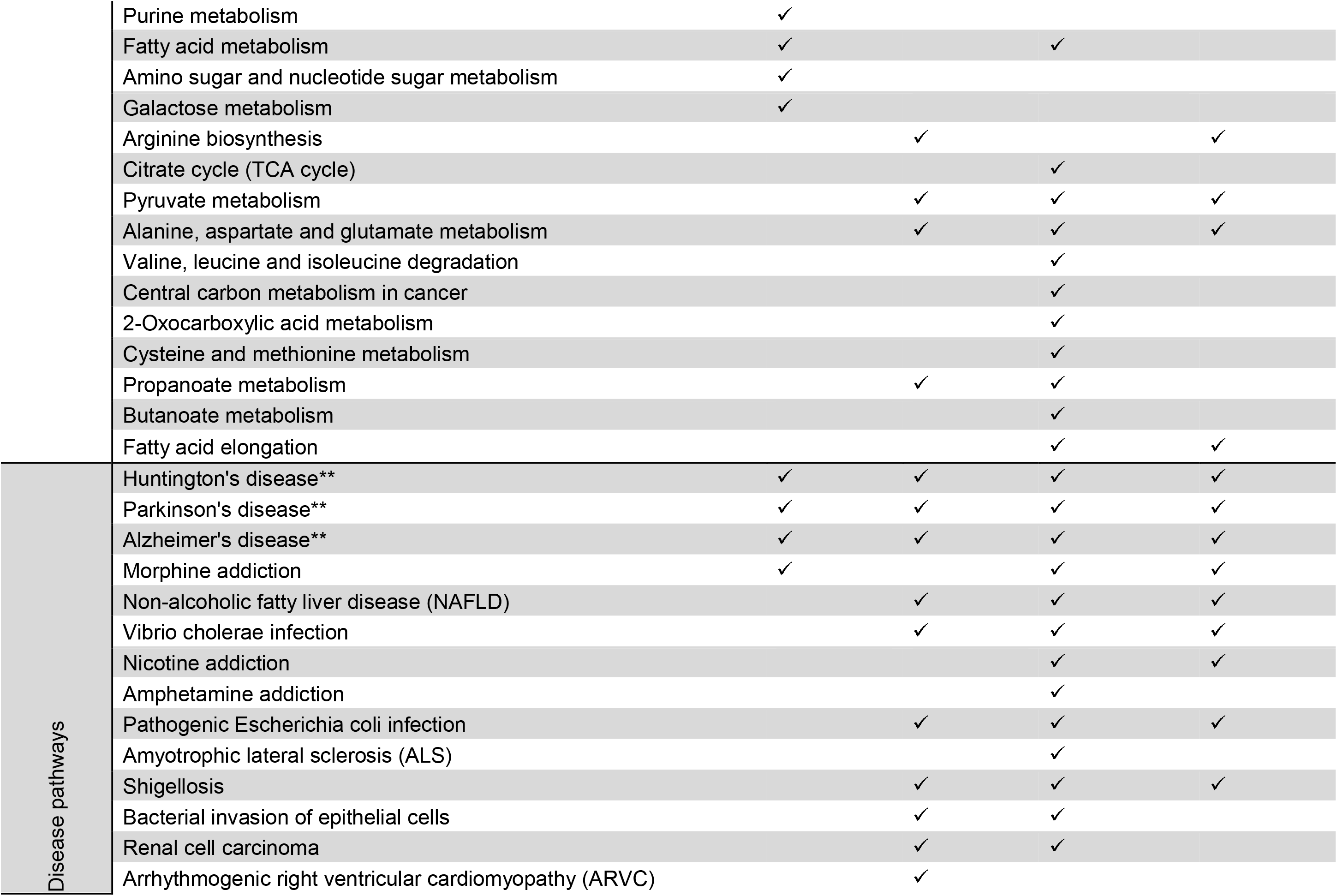
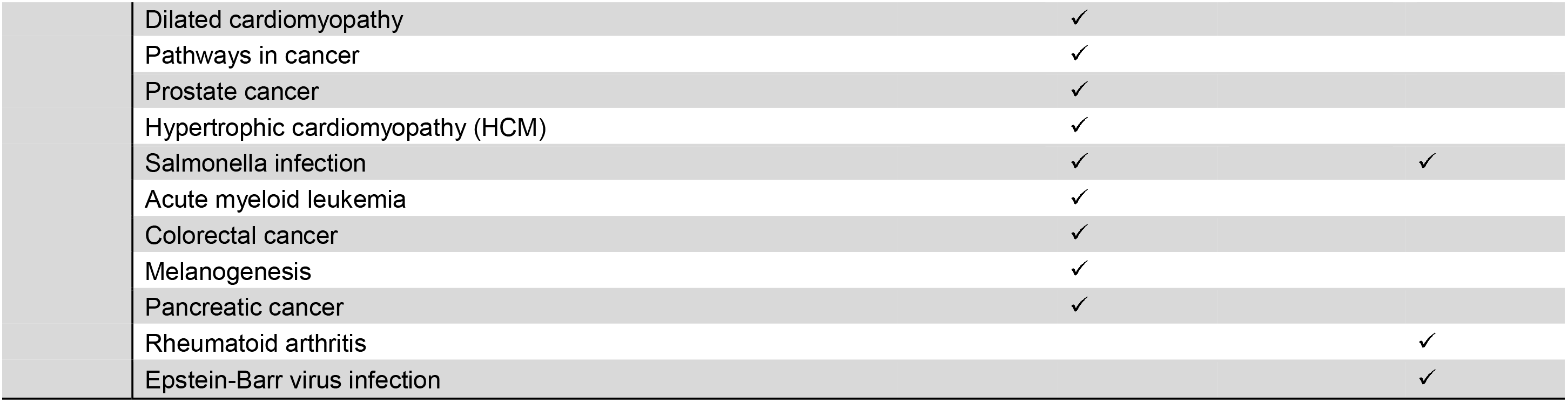
list of all pathways and their appearance in brain regions. Classification performed based on a function of pathways. Identical pathways were highlighted by two-star.

### 2-4 Network reconstruction

A network is a graph of nodes (genes/proteins) and edges (interactions) [32]. In this study, we reconstructed two networks based on knowledge-and data-driven approaches. First, a protein-protein interaction network (PPI) was constructed based on prior knowledge of signaling and metabolic pathways in different pathway databases (section 5-1). Then, a reverse-engineering approach was applied as a data-driven method (section 5-2).

#### 2-4-1 knowledge-driven network reconstruction

The Search Tool for the Retrieval of Interacting Genes/Proteins (STRING), a database for predicted protein-protein interactions at EMBL. It is clustering the extracted results from many protein-protein interactions databases, like Mint, BioGrid, etc. It also uses the information from KEGG pathways and Reactome to provide the best annotations for the interactions of one protein. We constructed a network by submitting DEG list to STRING database of protein-protein interaction to create Brain PPI Network in Alzheimer’s disease (BPNA) (Figure 6) representing the multi-regional network in the brain of Alzheimer’s patients. BPNA network was decomposed into five sub-networks to identify region-specific networks including BPNA_SEED_ (proteins of identical pathways which is described in section-4), BPNA_FRO_ (proteins of non-identical pathways of the Frontal region), BPNA_ENT,_ BPNA_HIPP_ and BPNA_TEM_ as is illustrated in figure 8 (Table S3).

#### 2-4-2 Data-driven network reconstruction

In this step, we have tried to reverse engineer a gene regulatory network (GRN). In this research, ARACNe (Algorithm for the Reconstruction of Accurate Cellular Networks) was used to construct a Gene Regulatory Network (GRN) for each of the brain regions [33, 34]. ARACNe network of each brain region was constructed based on the dataset with the best score in the MetaQC analysis. Mutual information(MI) threshold and Data Processing Inequality (DPI) tolerance set to 0.05 and 0%, respectively [35].

#### 2-5 Network Analysis

All constructed networks (PPI/GRN) were assessed for topological properties such as Betweenness centrality (B.C), closeness centrality (C.C), the degree of the nodes and topological coefficient (Table S4 and S5) [36, 37]. Additionally, Newman’s modularity (community) algorithm was applied for detection of the strength of division of the network into modules [38].

#### 2-6 Clustering and Correlation Analysis

Correlation between brain regions was performed using expression data of DEGs (section 2) from case (patients) samples of the GSE5281 dataset (best score in MetaQC test, refer to section 2) in R software by Kendall method and cor algorithm (stats package) [39]. To define effective genes in the correlation between regions, the pairwise correlation was performed using expression data of pooled DEGs from all samples of the GSE5281 dataset. Correlated gene pairs (the nodes) were connected by edges to create a correlation network. A number of edges and nodes of pairwise correlation were extracted and depicted by circos software [40] (Figure 4-a). Investigation of pairwise correlation between brain regions was performed to identify correlated genes (Correlation coefficient: 0.8 and p-value < 0.05). Pairwise correlations results hold as networks. Topological properties of all networks were analyzed and showed these networks were scale-free, so nodes with a degree above average called hubs. The average degree of nodes and frequencies of hubs in each network were calculated. A number of nodes/hubs were compared in each paired correlation network (Figure 4-b) and were compared with other pairs.

Clustering of selected genes between samples was performed to investigate related genes in brain regions based on their expression data of case samples using “dendextend” package [41] (Figure 5-b). The number of clusters was determined by “fviz dend” algorithm in this package (Figure S1). Results of dendrogram were shown in Table S6 and the complete results supplied in supplementary files (Figure S1). Clustered genes were enriched based on their class by Enrichr database (refer to section 3). For a better understanding of relationships between regions, clustering of brain regions was performed on data (Figure 5-a). Finally, expression of all probesets was applied to draw clustering dendrogram between regions (Figure 3).

**Figure 3-.**
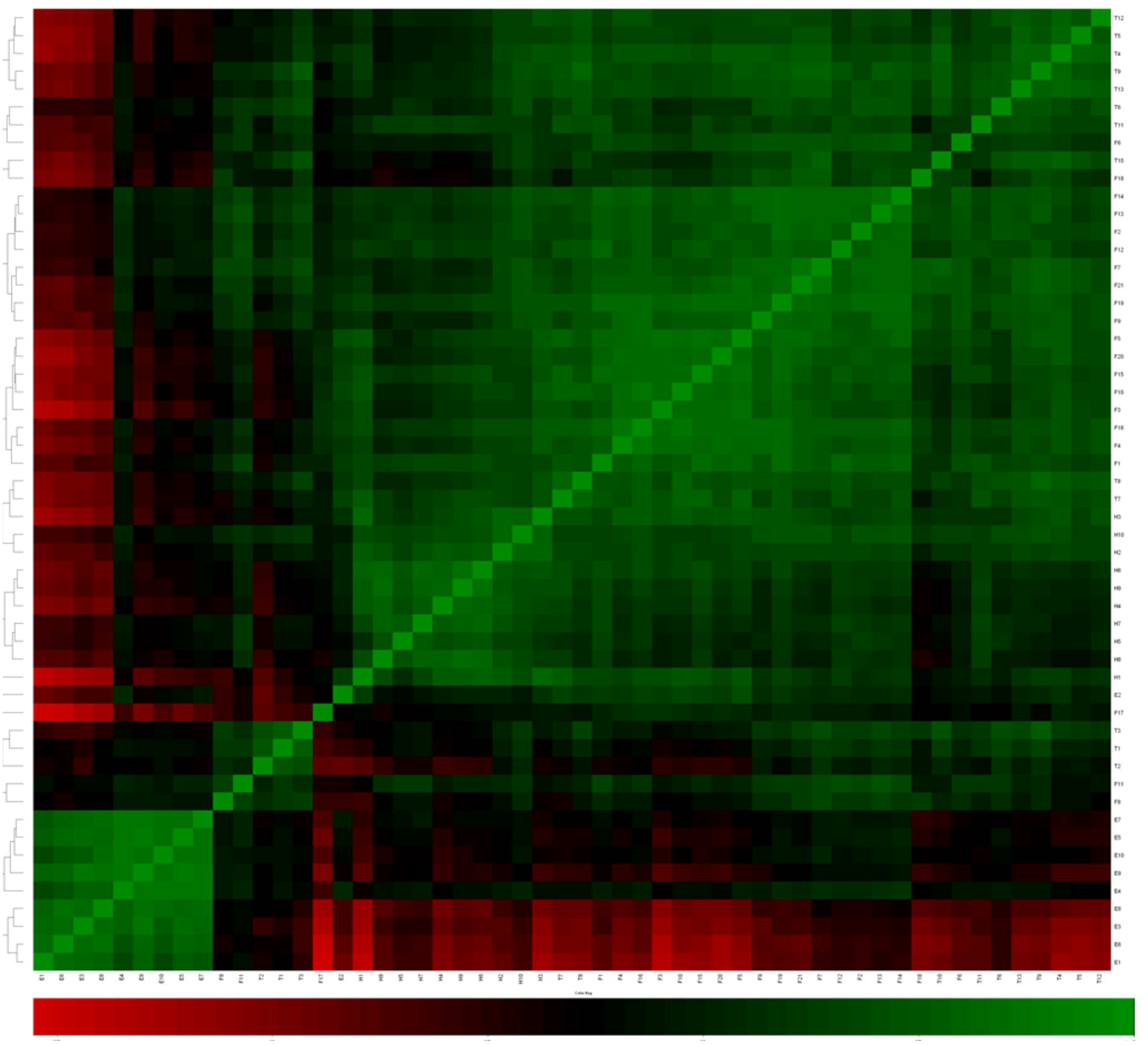
Correlation between brain regions. Entorhinal samples show a negative correlation with other regions of the brain in AD and were clustered in a distinct group.

**Figure 4-.**
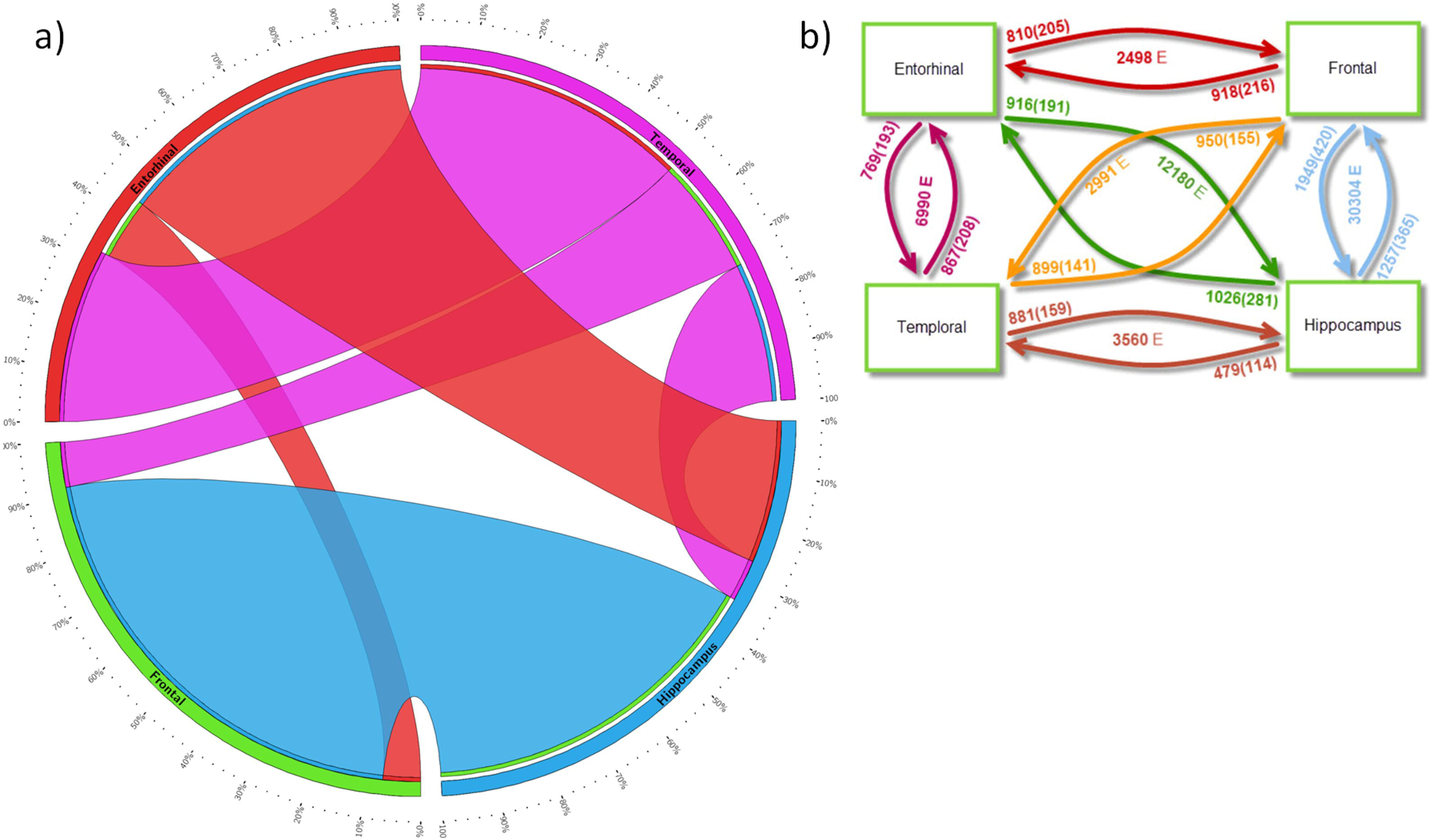
Pairwise correlation between brain regions. a) Schematic diagram based on a percentage of edges in Hippocampus-Frontal and Entorhinal-Temporal regions, respectively. b) Pairwise correlations showed by two arrows between brain regions. At the beginning of each arrow, a number of nodes in the adjacent region and its hubs nodes (within parenthesis) are illustrated. A Total number of edges are shown between paired arrows.

**Figure 5-.**
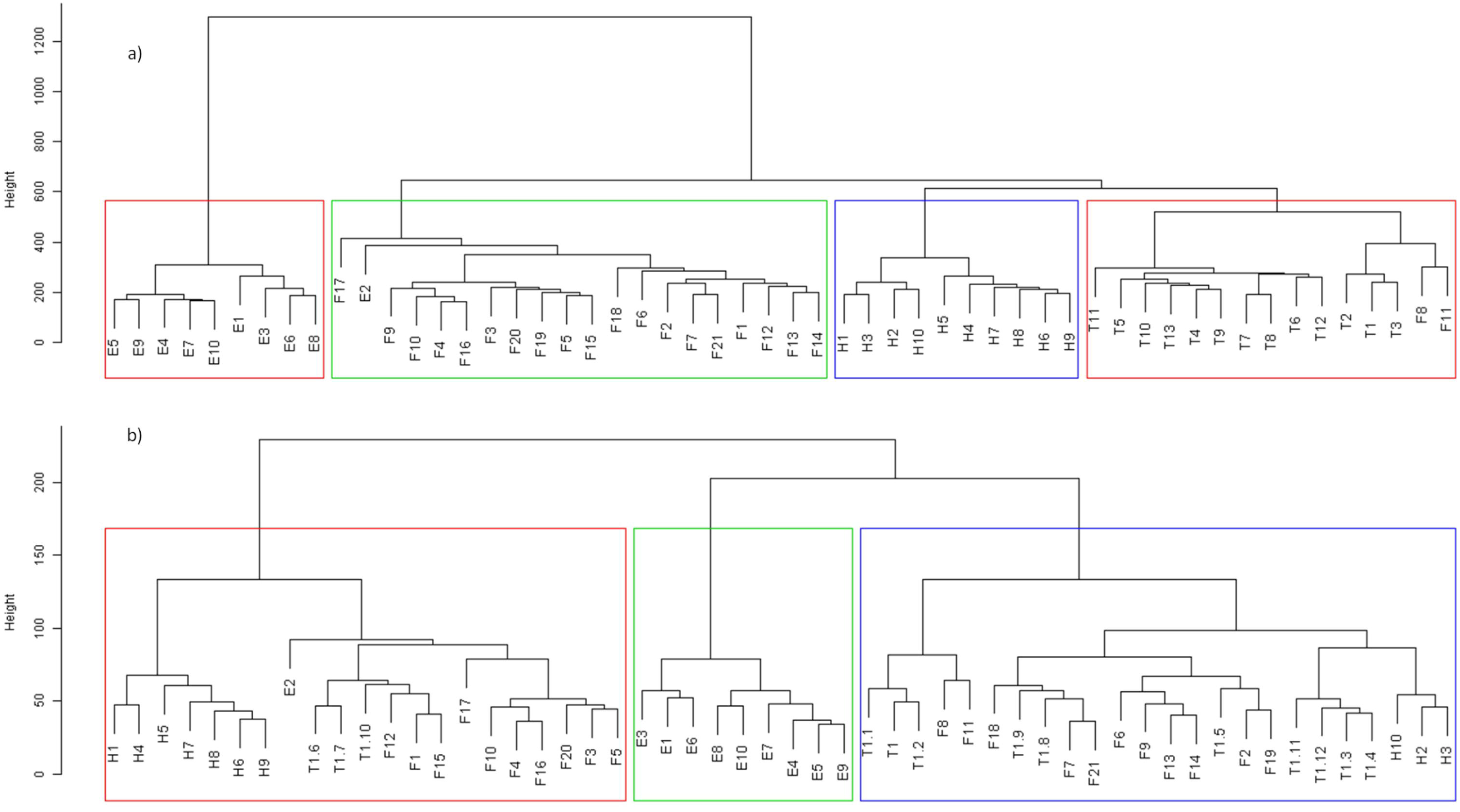
Dendrogram and heatmap of brain regions. a) clustering of brain regions based on all probeset expression data. Samples were clustered properly except for three of them including F8, F11, and E2. b) clustering of DEGs based on their expression showed that samples from Entorhinal region are classified in a distinct group. However, samples of other regions are classified into two groups and three subgroups except for F8, F11, and E2. Abbreviations are: H = Hippocampus, E = Entorhinal, F = Frontal and T = Temporal.

#### 2-7 Cross-validation

We used two methods of cross-validation including literature mining and reconstruction of gene regulatory network of ARACNe. First, differential and critically identified proteins in BPNA were used for literature mining (Table S7). Then, ARACNe network was constructed for each region of the brain. Afterward, results were compared between different methods. Nodes with a degree above average were selected as hubs. Also, Hubs were extracted and compared to intra-and inter-region correlation results.

## 3- Results

Despite the clinical progression and advanced studies in the field of Alzheimer`s disease (AD), its diagnosis and curation remain unclear. Many separated meta-analytical studies suggested the possibility of sharing a common feature among different AD brain regions [42–44]. Herein, we have tried horizontal and vertical data integration approach to unify heterogeneous microarray studies for several regions of the brain in AD patients and investigate their possible signaling and regulatory relationships. Also, a network approach was followed to investigate key effective genes in the initiation and progression of AD through different brain regions.

### 3-1 Meta-analysis and Gene expression analysis result

Among 110 transcriptomics brain studies in Alzheimer’s patients with about 2000 samples in overall, four brain regions including Entorhinal, Frontal, Hippocampus, and Temporal were selected for meta-analysis following quality assessment and just in Temporal region one sample was excluded (Figure 2 and Table 2). Afterward, DEGs in each brain region were identified (FDR≤10^−3^, fold change >|2|)”. Hippocampus region with 1438 genes presented the highest number of DEGs, and the Entorhinal region with 486 genes showed the lowest number of DEGs. Frontal and Temporal regions with 1131 and 998 genes ranked between Hippocampus and Entorhinal, respectively. Detailed results for meta-analysis of each brain region is provided in the supplementary file of Table S1.

By analyzing top 10% unique DEGs of each region, it was determined that each region has different functional pattern than another region. In Entorhinal region (with 307 unique DEGs), most of the top DEGs contribute in Metabolic pathways, such as amino acids, fatty acid, and carbon metabolism, but in Hippocampus (with 730 unique DEGs) oxidative phosphorylation, mitochondrial related pathways and ataxia were enriched. In Frontal region (with 407 unique DEGs) selected DEGs highlighted MAPK and RAS signaling pathways and neuropathy disease, however, in the Temporal region (with 419 unique DEGs) adherens junction was important pathway along with muscular dystrophy was the pathological term (Table S2).

### 3-2 Comparison of expression data among different brain regions

In the first step of correlation analysis, all genes in all samples of brain regions were clustered (Figure 3). Generally speaking, clustering of brain samples showed that Entorhinal has a negative correlation with other brain regions, while there are some positive and negative correlations between the samples from Frontal, Temporal and Hippocampus regions. To identify effective genes, pairwise correlation analysis was performed using selected genes in all samples (Figure 4-a). The resulting six pairwise correlation networks were constructed to identify shared edges between regions (Figure 4-a). Percent of similarity between different brain regions (number of nodes from one region/number of all nodes of that region) is illustrated in figure 4-a. This comparison showed that in each pairwise correlation, a region with a high average degree in nodes has a low frequency in hubs and may have a regulatory role over other regions. Figure 4-b is a summarized graph of the figure 4-a in which arrows show the pairwise correlation. Also, numbers indicate the nodes of each region that contribute in the pairwise correlation. Results showed that Entorhinal region has less number of hubs than other regions in each comparison.

The samples from different brain regions were classified based on; a-all expression data and b-DEGs. All four brain regions are clearly clustered when all genes are used for clustering (Figure. 5a). However, clustering of brain regions based selected DEGs showed different results (Figure 5b). Interestingly, almost all Entorhinal samples are clustered in one distinct group, and other samples of three other regions are co-clustered. Therefore, gene expression profile in Entorhinal shows a different pattern in comparison with other brain regions.

Additionally, DEGs were clustered in two distinct groups of up-regulated and down-regulated genes (Figure 6 and Table S6). Each cluster was used for gene enrichment analysis (Table S6). Neurodegenerative disease pathways (including AD, Huntington, and Parkinson) and oxidative phosphorylation pathways were enriched for the up-regulated cluster, but cell cycle and proteasome were enriched for the down-regulated cluster (Adjusted p-value < 0.05).

**Figure 6-.**
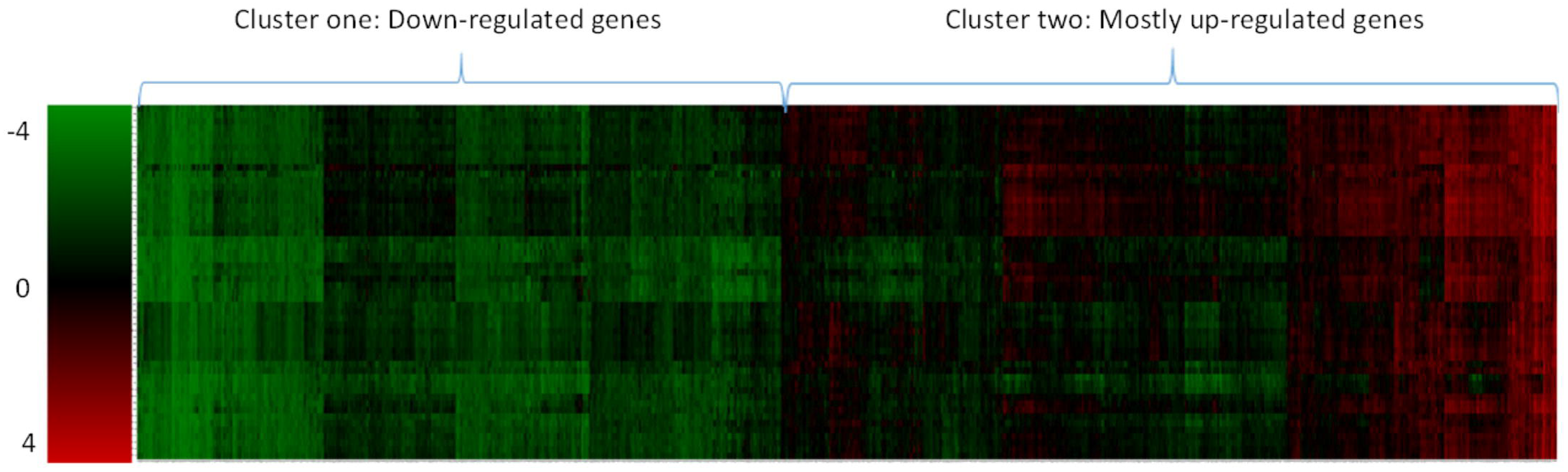
Heatmap of clustered DEGs in two distinct groups of up-regulated and down-regulated genes

### 3-3 Pathway enrichment and integration

After gene enrichment in Enrichr, Totally, about 97 KEGG’s pathways were identified for Entorhinal, Frontal, Hippocampus, and Temporal with eight overlapping pathways among these pathways (Table 2).

### 3-4 PPI network reconstruction and analysis

#### 3-4-1 knowledge-driven network analysis

Proteins of enriched pathways were merged and used to reconstruct BPNA PPI network in STRING database. BPNA was a scale-free network of 727 nodes and 8974 edges. The network comprised of eight modules. About 10% of BPNA nodes (75 nodes) belonged to KEGG Alzheimer’s disease pathway. The degree of about 78% of these nodes was above average. 66% of KEGG Alzheimer’s disease pathway was clustered in one module. Eight proteins were dysfunctional in all four regions including DNM1, GNG3, AP2M1, CDK5, ATP5G1, ALDOA, GPI, and TPI1. NDUFB1, NDUFA2, NDUFB4 and NDUFB11 and showed different behavior compared to other hub proteins in their closeness index. Interestingly, proteins of KEGG Alzheimer’s disease pathway tend to have a high degree.

To determine the effective pathways in each brain region and related to Alzheimer progression, BPNA network was decomposed into two sub-network groups based on eight identical pathways and non-identical pathways (enrichment results) of the brain region. Proteins of eight identical pathways were used to induce a subgraph called BPNA_seed_. The sub-network consisted of 196 DEGs from different brain regions (figure 7). Afterward, proteins from non-identical enriched pathways were used to induce four brain-region-specific sub-graphs including BPNA_FRO_ (208 proteins), BPNA_ENT_ (75 proteins), BPNA_HIPP_ (407 proteins) and BPNA_TEM_ (335 proteins).

**Figure 7-.**
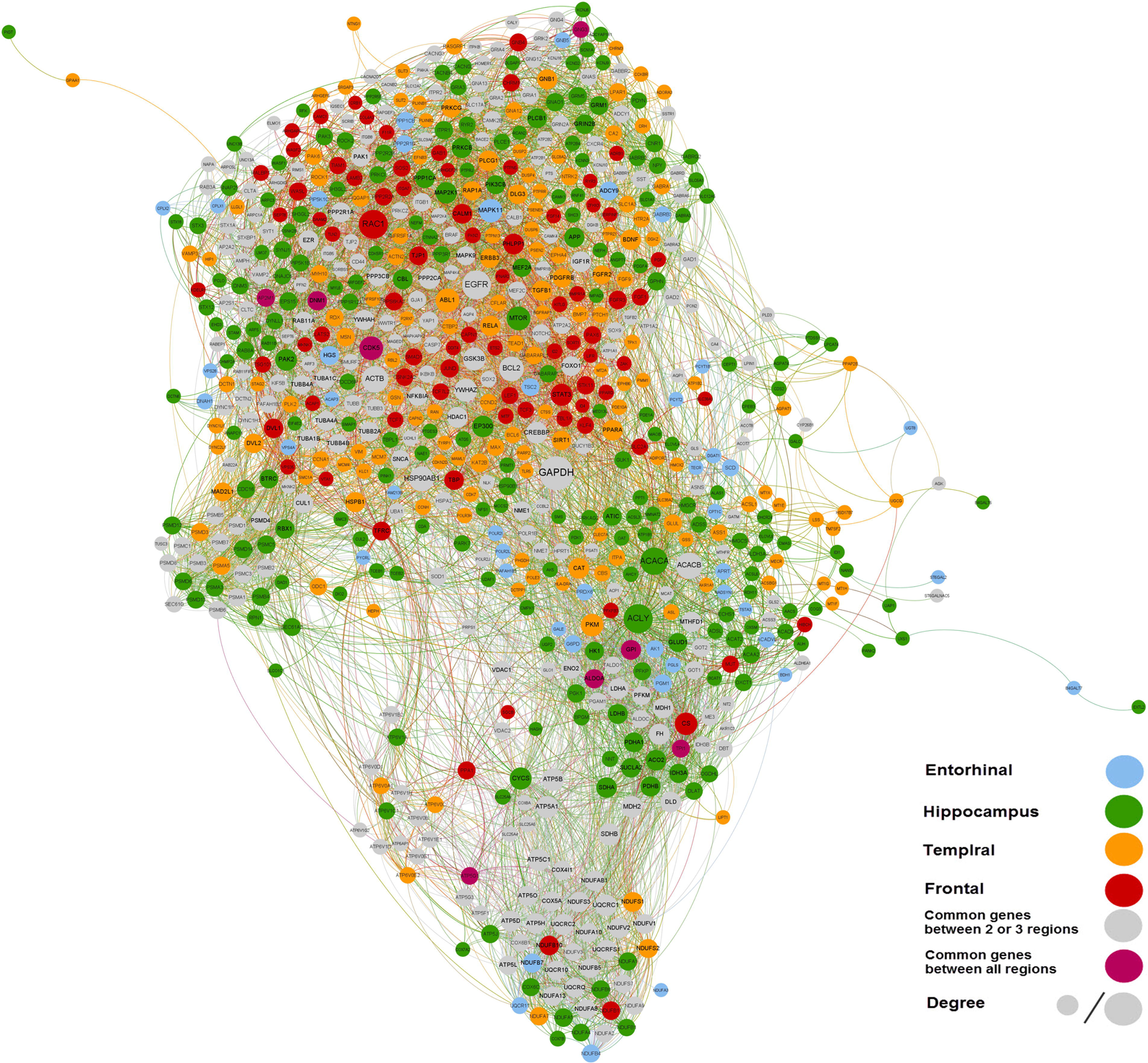
BPNA network contains 727 nodes. Most of the nodes belong to the hippocampus and Frontal regions. Eight proteins that are identical between all brain regions are shown in purple. GAPDH has the highest degree in the network.

The frequency of protein numbers in identical pathways for different brain regions in figure 8 indicate that the Hippocampus region includes the most proteins in each identical pathway, most of the non-identical pathways belonged to Hippocampus and Entorhinal includes the less number of pathways (Table 2).

**Figure 8-.**
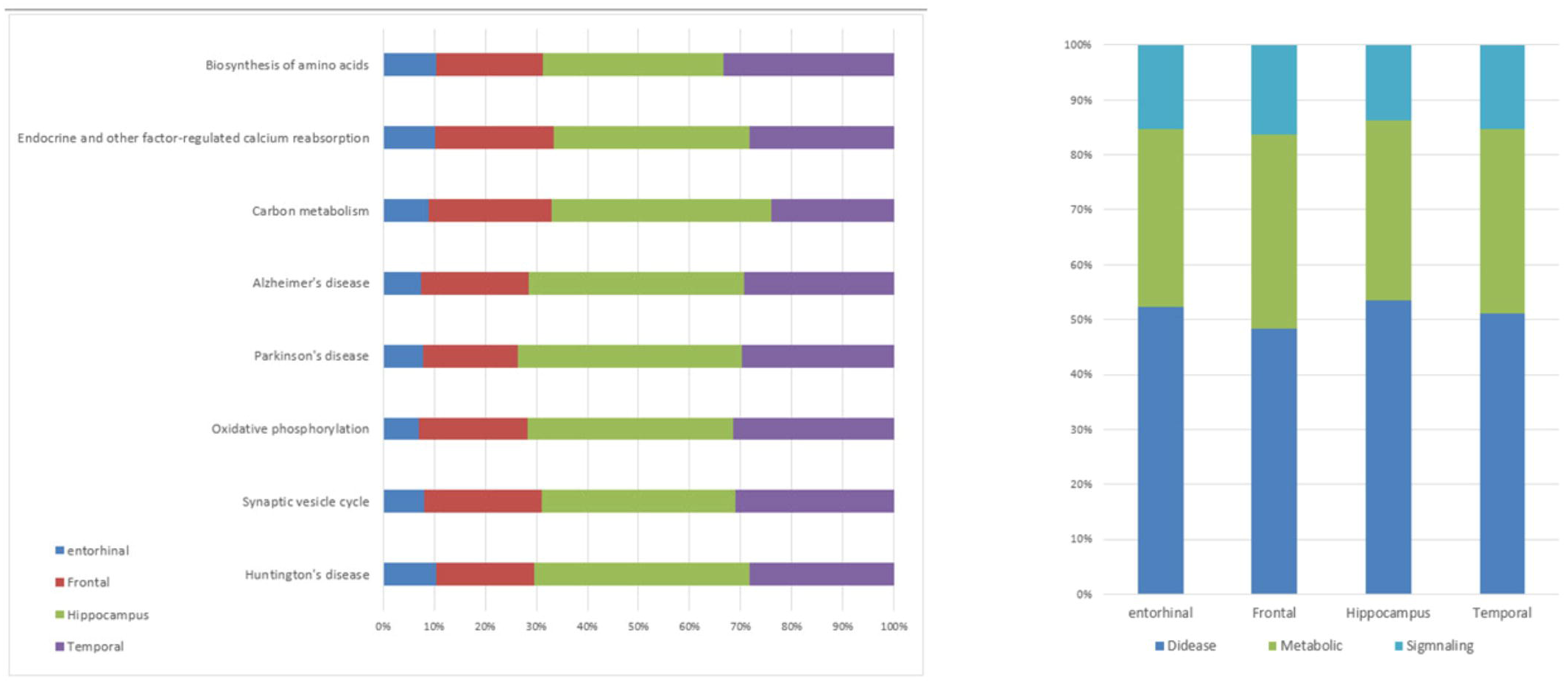
Comparison of genes in identical pathways in each brain region. a) Identical pathways contain three signaling, two metabolic and three pathological pathways. B) Comparison of frequencies of genes in BPNA sub-networks, based on their pathway class.

All decomposed networks constructed from BPNA are represented in figure 9 (proteins of all selected pathways). In BPNA_SEED_ network (Figure 9-a), disease pathways (24%), signaling pathways (16%) and metabolic pathways (12%) have the most number of specific proteins. Interestingly, 21 of 25 high degree proteins, belonged to KEGG Alzheimer’s disease pathway. UQCRC1, NDUFB1, NDUFB4, NDUFB11, NDUFA2, NDUFA5 and GAPDH were hub proteins of this brain region in AD patients. Interestingly, in this network GAPDH was special protein belonging to all three mentioned classes of pathways, also is a KEGG Alzheimer’s disease pathway protein and were in top ten lists of high degree proteins, first in high B.C and C.C and 9^th^ in low topological coefficient.

**Figure 9-.**
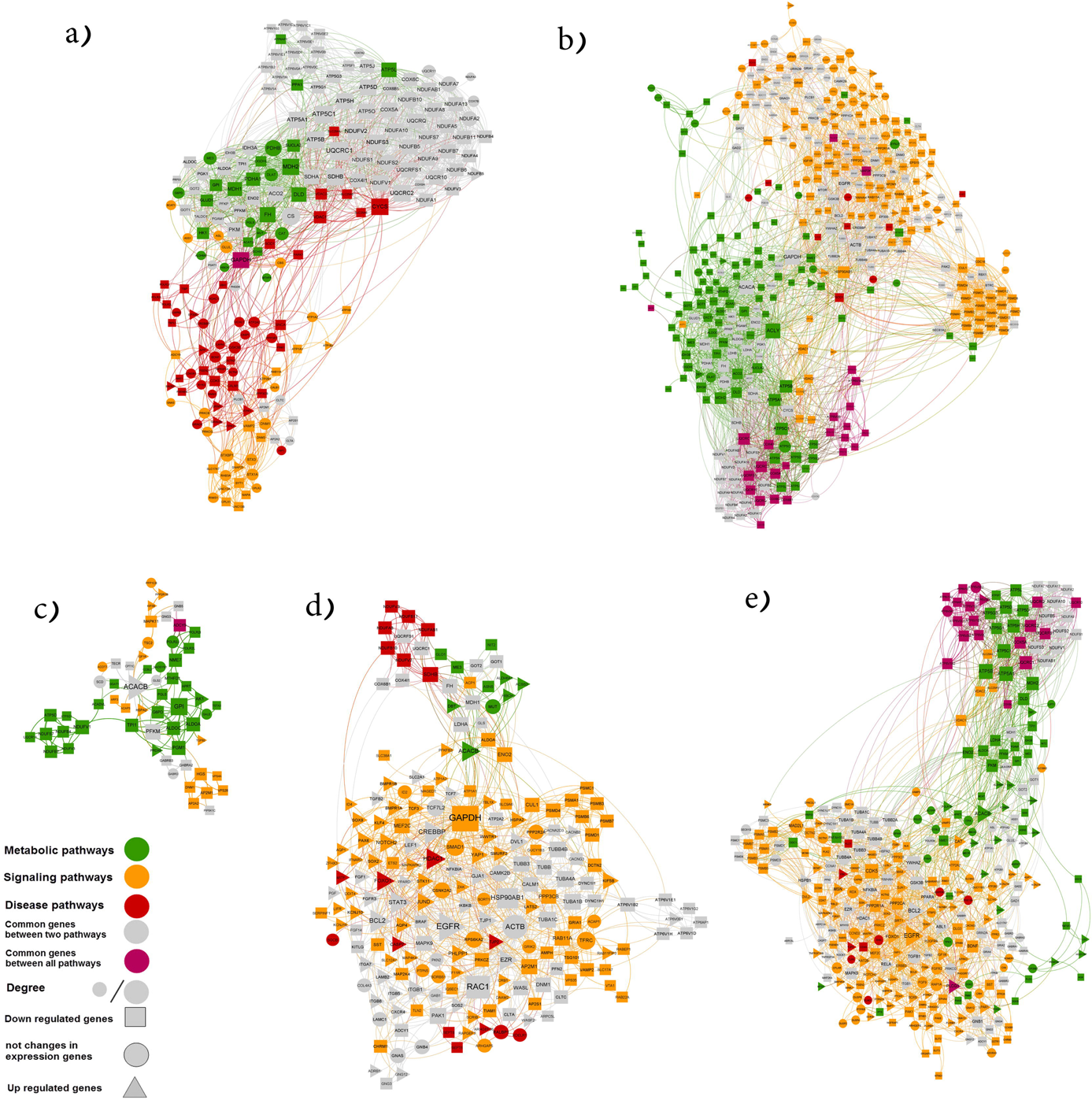
Decomposition of BPNA network into several subnetworks. All common genes between four different brains regions are colored in gray. Unique genes are colored based on the category of biological pathways (according to Table 2). The differences in gene expression based on metaomics analysis have been illustrated with different shapes (▴, • and ▪ for each of upregulated, downregulated and not-differential genes, respectively. (a) BPNA_SEED_ b) BPNA_ENT_ c) BPNA_FRO_ d) BPNA_HIPP_ e) BPNA_TEM_)

Figure 9 b-e is representing specific sub-graphs. In Entorhinal region (BPNA_ENT_), ACACB protein (expressed in both metabolic and signaling pathways) has the highest degree (Figure 9-c). In this subnetwork, NDUFB4, NDUFS7, NDUFB7, and ATP5D, were proteins that have a degree above average (Hub) but very low B.C and C.C.ADCY9 (the enzyme that catalyzes the formation of the cyclic AMP from ATP) was the only protein that contributed in all three classes of pathways.

In BPNA_FRO_ GAPDH (a protein of KEGG Alzheimer’s disease pathway) had the highest degree, (Figure 9-b). BPNA_TEM_ showed different characterization. This network has 20 proteins (6% of all nodes of this network) that contributed in all signaling, metabolic and disease pathways and interestingly almost all of them modularized in oxidative phosphorylation pathway (based on KEGG Mapper analysis). These proteins include UQCRFS1, UQCR10, UQCRQ, UQCRC2, COX5A, UQCRC1, COX8A, ATP6V1H, ATP6V1B2, ATP6V0A1, ATP6V0C, ATP6V1E1, ATP6V0D1, ATP6V0E2, ATP6V0B, ATP6V0E1, ATP6V1G2, ATP6AP1, ATP6V1D, and PLCG1. First seven proteins also contribute to KEGG Alzheimer’s disease pathway. EGFR included the highest degree in BPNA_TEM_ (Figure 9-e).

Finally, BPNA_HIPP_ contained 391 nodes and was the largest sub-network. ACLY had the highest degree in BPNA_HIPP_ network (Figure 9-d).

#### 3-4-2 Data-driven network reconstruction: ARACNe

Based on the MetaQC analysis, GSE5281 data series were selected for ARACNe analysis in all four brain regions. The resulted ARACNe networks included 5380 nodes and 28795 edges in Entorhinal, 4993 nodes and 12835 edges in Frontal, 4732 nodes and 29044 edges in Hippocampus and 5337 nodes and 42547 edges in the Temporal region. To have a glance at resulted networks, top 10 genes in degree and their first neighbors were extracted, and new networks were enriched, separately. Pathways with adjusted p-value < 0.05 were selected and shown in Table 3. Entorhinal regions showed the minimum and different pathways than other regions, indeed just two pathways of Entorhinal shared with another region (Temporal) and highlighted in Table 3. But three remained regions have four shared pathways including Oxidative phosphorylation, Synaptic vesicle cycle, epithelial cell signaling in Helicobacter pylori infection and finally Vibrio cholerae infection.

**Table 3-.** Comparison between selected pathways that are resulted from the enrichment of top 10 genes in degree and their neighbors of GRN networks.

|  | GRN<sub>TEM</sub> | GRN<sub>HIPPO</sub> | GRN<sub>FRO</sub> | GRN<sub>ENT</sub> |
| --- | --- | --- | --- | --- |
| Selected pathways | Oxidative phosphorylation** | Oxidative phosphorylation** | Oxidative phosphorylation** | GABAergic synapse<sup>#</sup> |
|  | Synaptic vesicle cycle** | Synaptic vesicle cycle** | Synaptic vesicle cycle** | Glutamatergic synapse<sup>#</sup> |
|  | Epithelial cell signaling in Helicobacter pylori infection** | Epithelial cell signaling in Helicobacter pylori infection** | Epithelial cell signaling in Helicobacter pylori infection** | Ribosome biogenesis in eukaryotes |
|  | Vibrio cholerae infection** | Vibrio cholerae infection** | Vibrio cholerae infection** | Serotonergic synapse |
|  | Cardiac muscle contraction | Cardiac muscle contraction | Alzheimer's disease | Aminoacyl-tRNA biosynthesis |
|  | Retrograde endocannabinoid signaling | Retrograde endocannabinoid signaling | Parkinson's disease | Central carbon metabolism in cancer |
|  | Carbon metabolism | Epstein-Barr virus infection | Epstein-Barr virus infection | Citrate cycle (TCA cycle) |
|  | Glutamatergic synapse | Collecting duct acid secretion | Collecting duct acid secretion | Morphine addiction |
|  | GABAergic synapse | Rheumatoid arthritis | Rheumatoid arthritis | Regulation of autophagy |
|  | Alzheimer's disease | Metabolic pathways | Metabolic pathways |  |
|  | Amyotrophic lateral sclerosis (ALS) | Proteasome | Proteasome |  |
|  | MAPK signaling pathway | Endocytosis | Endocytosis |  |
|  | Alanine, aspartate and glutamate metabolism | Cytosolic DNA-sensing pathway | Non-alcoholic fatty liver disease (NAFLD) |  |
|  | Phenylalanine, tyrosine and tryptophan biosynthesis | Fatty acid elongation | Salmonella infection |  |
|  | Glycolysis / Gluconeogenesis | RNA polymerase | Huntington's disease |  |
|  | Nicotine addiction | TNF signaling pathway | Phagosome |  |
|  | Arginine biosynthesis | Cholinergic synapse | ErbB signaling pathway |  |
|  | Biosynthesis of amino acids | Fatty acid metabolism | Colorectal cancer |  |
|  | Cysteine and methionine metabolism | Chemokine signaling pathway |  |  |
|  | Huntington's disease |  |  |  |
|  | Phagosome |  |  |  |

### 3-5 Cross-validation

All differentially expressed genes during this meta-analysis are available in Table S7. We have reported 100 Up and 486 down-regulated genes in Alzheimer disease in all brain regions completely that more than 75% of our reported in each group were validated. Summary of Cross-validation statistic is shown in Table 4.

**Table 4-.** Summary of Cross-validation statistics. Numbers of up or down-regulated genes per brain region and percent of their validity in literature were shown.

| **Regions** | **Up-regulated genes**<br>**NO** | **Validity percent** | **Down-regulated**<br>**genes NO** | **Validity percent** |
| --- | --- | --- | --- | --- |
| **Hippocampus** | 25 | 84 | 229 | 98 |
| **Temporal** | 29 | 76 | 137 | 99 |
| **Frontal** | 41 | 93 | 79 | 94 |
| **Entorhinal** | 5 | 100 | 41 | 95 |

## 4- Discussion

Relating global expression pattern of multiple genes in AD to the different regions of the brain and deciphering AD progression in these regions is a serious part of the research because AD is one of the most common neurodegenerative disorders. Some related studies on brain regions of Alzheimer patients were accomplished previously. Ray *et al*. by analyzing microarray data of brain regions showed that middle temporal gyrus is a good choice for detecting early AD pathogenesis [43]. Wang *et al*. extract sample from 19 cortical regions of 125 individuals and after defining DEGs and ranking brain region based on relevance to AD, construct co-expression networks for each brain region. Their analyses identified temporal lobe gyri as earliest gene expression abnormalities region [45].

Our meta-analysis results showed that Entorhinal region has the minimum number of DEGs with respect to other regions. Also in some analysis results, Entorhinal showed consistently unique behavior compared to other regions: A) In correlation analysis: displayed negative correlation with other brain regions (Figure 3), B) In clustering of all DEGs (Figure 5-b), Entorhinal samples were classified in a separate from samples of other regions C) In Gene set enrichment analysis, Entorhinal region contained non-identical pathways with metabolic pathways owning the highest number of pathways, while in other regions, the highest number of pathways were related to signaling pathways, Entorhinal had also few hubs with high average degree in each pairwise correlation (Table 2) and finally D) In pathway enrichment analysis on selected genes in GRN analysis, list of Entorhinal pathways were different compared to other regions (Table 3). Interestingly, there are several integrative observations indicating the beginning of AD from entorhinal [46–49]. Mapping of histology and imaging-based data (MRI and PET) in different studies have revealed the critical and initial function of the entorhinal region in AD pathogenesis. We have also observed different expression pattern for entorhinal (more hubs and inverse pairwise correlation) in comparison to other studied regions.

The results of meta-analysis also indicated the highest number of DEGs in Hippocampus. Pairwise and correlation gene expression analysis between regions of brain w showed that Entorhinal region has different manner with respect to other regions and control another part of the brain (see above). Reviewing clinical case studies on the AD patient whose disease progression has been monitored using brain imaging techniques demonstrated that brain atrophy initiated from the limbic lobe (Entorhinal and Hippocampus regions) to Temporal and will reach to Frontal region [50]. Moreover, the Hippocampus and Entorhinal atrophy is a clinically relevant AD diagnostic measure in patients [51]. Interestingly, our results impress on the importance of these two regions. Enrichment results for top 10% differential genes in each brain region indicate their different functioning in which hippocampus, frontal and temporal regions were enriched in pathological terms including ataxia, neuropathy and muscular dystrophy, respectively [52–54].

A clear crosstalk was observed between various brain regions in AD patients so related pathways-some of which are shared among regions-were functioning simultaneously. Also, two distinct classes of pathways were over-(neurodegenerative diseases and related pathways) and under-(cell cycle and proteasome) represented. Because of limitation in data sources, results of this research were not ideal, and by sampling from all brain regions of AD patients, can perform completely. Altogether, application meta-analysis in connects to complete sampling resulted in identifying beginning and progression factors of AD.

We observed that GAPDH had the highest node degree protein in Hippocampus and frontal regions, in addition, this protein was connected to proteins in all brain regions. GAPDH isoforms have been found in the brains of AD patients [55]. The eight proteins which were dysfunctional in all four regions were directly involved in the pathogenesis of AD. ALDOA, GPI, and TPI1 are involved in inflammatory pathways in AD via induction of inflammation [56], activation of β-secretase [57], and nitrotyrosination of proteins, respectively. AP2M1 and DNM1 contribute to the degradation of β-amyloids and abnormal mitochondrial dynamics which lead to impaired synaptic degeneration and signaling, respectively [58–60]. GNG3 another member of this list encodes gamma subunit of Guanine nucleotide-binding protein G(I)/G(S)/G(O) which is required for GTPase and is its dysregulation is involved in the initiation of AD [61]. Cdk5 and ATP5G1 are involved in mitochondrial dysfunction in AD. CDK5 is one of the most studied and critical proteins in AD pathogenesis involved in neuronal migration, differentiation, synaptic functioning and memory consolation. Its hyperactivity and deregulation cause many of pathological outcomes in AD [62]. Also, decreased activity of ATP5G1 (ATP synthase) along with its partners in respiratory chain is reported in AD [63].

In all depicted networks, proteins involved in AD pathogenesis tend to contribute in one module, showing high relation and communication (such as protein-protein interactions) between them. Also, most of the AD genes were hubs in mentioned networks that can be related their active roles in samples. Additionally, a wide range of effective proteins was identified which are belonging to glycolysis (GADPH), mitochondrial electron chain (ATP5C1) or folding of proteins (HSPA8), the main effective factors in AD progression. Combining proper microarray data in this meta-analysis, and their comparison concluded to a noteworthy clustering.

## 5- Conclusion

In this study, we investigated the pattern of damage flow through multiple regions of the brain in AD using meta-analysis techniques. Moreover, the result of this study is consistent with former findings on protein expression pattern in the network of neurodegenerative diseases with special focus on AD. By linking the sequence of disease progression through multiple brain regions with DEGs observed in each brain area, the key role of underlying genes associated with initiation and progression of the disease. Besides, we showed meta-analysis techniques to be reliable in studying effective factors in disease progressing by creating an overall scope of operating disease factors. Altogether, we have identified that entorhinal shows different expression profiling than other three studies regions.

